# Recursive Prefix-Free Parsing for Building Big BWTs

**DOI:** 10.1101/2023.01.18.524557

**Authors:** Marco Oliva, Travis Gagie, Christina Boucher

**Affiliations:** Department of Computer and Information Science and Engineering, Herbert Wertheim College of Engineering, University of Florida, Gainesville, FL, USA; Faculty of Computer Science, Dalhousie University, Halifax, NS, Canada

**Author notes:** MO, and CB are funded by the National Science Foundation NSF SCH: INT (Grant No. 2013998), NSF IIBR (Grant No. 2029552), and National Institutes of Health (NIH) NIAID (Grant No. HG011392 and R01AI141810). TG is funded by NSF IIBR (Grant No. 2029552), NIH NIAID (Grant No. HG011392), and NSERC Discovery Grant RGPIN-07185-2020.

## Abstract

Prefix-free parsing is useful for a wide variety of purposes including building the BWT, constructing the suffix array, and supporting compressed suffix tree operations. This linear-time algorithm uses a rolling hash to break an input string into substrings, where the resulting set of unique substrings has the property that none of the substrings’ suffixes (of more than a certain length) is a proper prefix of any of the other substrings’ suffixes. Hence, the name prefix-free parsing. This set of unique substrings is referred to as the *dictionary*. The *parse* is the ordered list of dictionary strings that defines the input string. Prior empirical results demonstrated the size of the parse is more burdensome than the size of the dictionary for large, repetitive inputs. Hence, the question arises as to how the size of the parse can scale satisfactorily with the input. Here, we describe our algorithm, *recursive prefix-free parsing*, which accomplishes this by computing the prefix-free parse of the parse produced by prefix-free parsing an input string. Although conceptually simple, building the BWT from the parse-of-the-parse and the dictionaries is significantly more challenging. We solve and implement this problem. Our experimental results show that recursive prefix-free parsing is extremely effective in reducing the memory needed to build the run-length encoded BWT of the input. Our implementation is open source and available at https://github.com/marco-oliva/r-pfbwt.

## Introduction

The Burrows-Wheeler transform (BWT) [1] is fundamental to countless applications in bioinformatics, with one of the most notable applications being read alignment. If we consider all possible rotations of a given input text *T* sorted lexicographically, the *BWT matrix* is the matrix of all these rotations. The BWT array (denoted as BWT) is the last column of these rotations. Two powerful proprieties of the BWT are that it can be constructed from *T* in O(|*T*|) space and time, and it supports queries of the form: find the number of occurrences of the longest match between a query string *P* and *T* in *T*. These two properties stem from a fundamental property of the BWT called LF-mapping, which connects the last column of the BWT matrix (i.e. the BWT) with the first column of the BWT matrix. and the property that the BWT is reversible in O(|*T*|) -space and -time, Sequence alignment methods—such as BWA [2] and Bowtie [3]—build the BWT (and the suffix array) for a reference genome(s) which is then used to find short, exact alignments between each sequence read and the genome(s); these alignments are then extended using dynamic programming. Given the importance of the BWT to bioinformatics research, there have been many approaches developed and implemented for building the BWT for large datasets [4, 5]. Yet, construction algorithms still have a way to go before they can be used to construct the index for terabytes of genomic data. To date, the current largest BWT was constructed for approximately 1.7 terabytes of data via splitting the data into smaller parts, building the BWT of these parts, and then merging the obtained BWTs succinctly [5].

One of the most recent methods for building the BWT is `Big-BWT`, which first preprocesses the input using an algorithm called *prefix-free parsing* [6]. This preprocessing algorithm takes as input a string *T*, and two integers *w* and *p*, and then uses a rolling hash to find all *w*-length substrings in *T* that have hash value equivalent to 0 mod *p*, which are referred to as *trigger strings*. The set of trigger strings is used to define all unique substrings found in *T*, such that those strings start and end with a trigger string and contain no other trigger string. The lexicographically sorted set of said substrings is referred to as the *dictionary* (denoted as D). Given D we can define the list of occurrences of the elements of D in *T*, this list is referred to as the *parse* (denoted as P). Boucher et al. [6] showed that the BWT of *T* can be constructed from the dictionary D and parse P alone, without storing the input text *T*. This greatly improved the construction of the BWT, which currently remains unsurpassed. However, in our experience the space usage of the parse grows linearly while the size of the dictionary grows significantly slower.

In this paper, we address this final hurdle in building the BWT for large genomic databases: reducing the computational footprint of handling P. Our solution is to run prefix-free parsing on the parse obtained from running prefix-free parsing on the input text *T*. Hence, we refer to our algorithm as *recursive prefix-free parsing* because it generates the prefix-free parse of the prefix-free parse of the input. We denote the output of recursive prefix-free parsing as P_2_ and D_2_. This brings down the size of the parse by one order of magnitude, however building the BWT of T in space proportional to P_2_, D_2_, and D_1_ creates several algorithmic challenges. We implement our new approach for building the BWT, which we refer to as `r-pfbwt`. We show that `r-pfbwt` is 2.2 times faster, and requires 2.3 times less memory than `Big-BWT` on 2400 copies of Chromosome 19. Moreover, the empirical results demonstrate that the performance gains increase as the dataset gets larger.

## Preliminaries

### Basic definitions

A string *T* is a finite sequence of symbols *T* = *T* [1..*n*] = *T* [1] … *T* [*n*] over an alphabet Σ = {*c*_1_, …, *c*_σ_} whose symbols can be unambiguously ordered. We denote by *ε* the empty string, and the length of *T* as |*T*|. We denote as *c*^*k*^ as the string formed by the character *c* repeated *k* times.

We denote by *T* [*i*..*j*] the substring *T* [*i*] … *T* [*j*] of *T* starting in position *i* and ending in position *j*, with *T* [*i*..*j*] = *ε* if *i* > *j*. For a string *T* and 1 ≤ *i* ≤ *n, T* [1..*i*] is called the *i*-th prefix of *T*, and *T* [*i*..*n*] is called the *i*-th suffix of *T*. We call a prefix *T* [1..*i*] of *T* a *proper prefix* if 1 ≤ *i* < *n*. Similarly, we call a suffix *T* [*i*..*n*] of *T* a *proper suffix* if 1 < *i* ≤ *n*. Given a set of strings 𝒮, 𝒮 is *prefix-free* if no string in *S* is a prefix of another string in 𝒮.

We denote by ≺ the lexicographic order: for two strings *T*_2_[1..*m*] and *T*_1_[1..*n*], *T*_2_ ≺ *T*_1_ if *T*_2_ is a proper prefix of *T*_1_, or there exists an index 1 ≤ *i* ≤ *n, m* such that *T*_2_[1..*i* − 1] = *T*_1_[1..*i* − 1] and *T*_2_[*i*] < *T*_1_[*i*]. Symmetrically we denote by ≺_colex_ the co-lexicographic order, defined to be the lexicographic order obtained by reading *T*_1_ and *T*_2_ from right to left instead that from left to right.

### Suffix Array and Burrows-Wheeler Transform

Given a string *T* [1..*n*], the *suffix array* [7], denoted by SA_*T*_, is the permutation of *{*1, …, *n}* such that *T* [SA_*T*_ [*i*]..*n*] is the *i*-th lexicographically smallest suffix of *T*. We refer to SA_*T*_ as SA when it is clear from the context.

The Burrows-Wheeler transform of a string *T* [1..*n*], denoted by BWT_*T*_, is a reversie permutation of the characters in *T* [1]. If we assume *T* is terminated by a special symbol $ that is lexicographically smaller than any other symbol in Σ, we can define BWT_*T*_ [*i*] = *T* [SA_*T*_ [*i*] − 1 mod *n*] for all *i* = 1, …, *n*.

### Overview of Prefix-Free Parsing

Prefix-free parsing (PFP) takes as input a string *T* of length *n*, and two integers greater than 1, which we denote as *w* and *p*. It produces a parse of *T* consisting of overlapping phrases, where each unique phrase is stored in a dictionary. We denote the dictionary as D and the parse as P. We refer to prefix-free parse of *T* as PFP(*T*). As the name suggests, the parse produced by PFP has the property that none of the suffixes of length greater than *w* of the phrases in D is a prefix of any other. We formalize this property through the following lemma.

#### Lemma 1

([8]). *Given a string T and its prefix-free parse PFP* (*T*), *consisting of the dictionary* D *and the parse* P, *the set* 𝒮 *of distinct proper phrase suffixes of length at least w of the phrases in* D *is a prefix-free set*.

The first step of PFP is to append *w* copies of # to *T*, where # is a special symbol lexicographically smaller than any element in Σ, and *T* does not contain *w* copies of #. For the sake of the explanation, we consider the string *T* ′ = #^*w*^*T* #^*w*1^. Next, we characterize the set of trigger strings E, which define the parse of *T*. Given a parameter *p*, we construct the set of trigger strings by computing the Karp-Rabin hash, *H*_*p*_(*t*), of substrings of length *w* by sliding a window of length *w* over *T* ′ = #^*w*^*T* #^*w*^, and letting E be the set of substrings *t* = *T* ′[*s*..*s* + *w* − 1], where *H*_*p*_(*t*) ≡ 0 or *t* = #^*w*^. This set E will be used to parse #^*w*^*T* #^*w*^.

Next, we define the dictionary D of PFP. Given a string *T* and a set of trigger strings E, we let D = {*d*_1_, …, *d*_*m*_}, where for each *d*_*i*_ ∈ *D*: *d*_*i*_ is a substring of #^*w*^*T* #^*w*^, exactly one proper prefix of *d*_*i*_ is contained in E, exactly one proper suffix of *d*_*i*_ is contained in E, and no other substring of *d*_*i*_ is contained E. Hence, we can build D by scanning #^*w*^*T* #^*w*^ to find all occurrences of the trigger strings in E, and adding to D each substring of #^*w*^*T* #^*w*^ that starts at the beginning of one occurrence of a trigger string and ends at the end of the next one. Lastly, the dictionary is sorted lexicographically. Given the sorted dictionary D and input string *T*, we can easily parse *T* into phrases from D with consecutive phrases overlapping by *w* characters. This defines the parse P as an array of indexes of the phrases in D. We note that *T* can then be reconstructed from D and P alone. We illustrate PFP using a small example. We let *w* = 2 and

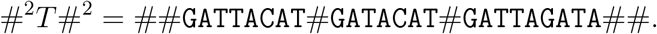

Now, we assume there exists a Karp-Rabin hash that define the set of trigger strings to be *{*`AC`, `AG`, `T`#, ##*}*. It follows that the dictionary D is equal to

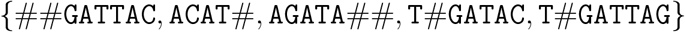

and the parse P to be [1, 2, 4, 2, 5, 3]. PFP can be used as a preprocessing step to build data structures such as the BWT and the SA.

In the next section, we will review how to compute the BWT of a string *T* using D and P.

### *Overview of* Big-BWT

Given the prefix-free parsing of a string *T*, we show how to build the BWT of *T* using |*PFP* (*T*) | space, i.e., |D| + |P| space, where |D| is the sum of the length of its phrases and P the number of elements in it. This is referred to as the `Big-BWT` algorithm. We will use the following properties of PFP that were first introduced [8], and outlined in the following lemmas. Moreover, we will refer to them later when describing our recursive algorithm.

To determine the relative order of the characters in the BWT_*T*_ —and hence, the relative lexicographic order of the suffixes following those two characters in *T* —we start by considering the case in which two characters are followed in *T* by two distinct proper phrase suffixes *α, β* ∈ 𝒮. From Lemma 1 it follows

#### Corollary 1

([8]). *If two characters T* [*i*] *and T* [*j*] *are followed by different phrase suffixes α and β, where* |*α*| ≥ *w and* |*β*| ≥ *w, then T* [*i*] *precedes T* [*j*] *in the BWT of T if and only if α* ≺ *β*.

In other words, for some of the characters in BWT_*T*_, it is sufficient to only consider the proper phrase suffixes which follows them in *T* to break the ambiguity. When this is not enough, we need the information contained in the parse.

#### Lemma 2

([8]). *Let t and t*′ *be two suffixes of T that begin with the same proper phrase suffix α, and let q and q*′ *be the suffixes of P that have the last w characters of those occurrences of α and the remainders of t and t*′. *If t* ≺ *t*′ *then q* ≺ *q*′.

Next, we give some intuition on how these two lemmas are used to compute the BWT. For each distinct proper phrase suffix *α* ∈ 𝒮 of length at least *w*, we store the range in the BWT containing the characters immediately preceding in the string *T* the occurrences of *α*. The starting position of the range for *α* is the sum of the frequencies in *T* (or P) of the proper phrase suffixes of length at least *w* that are lexicographically less than *α*. The length of the range is the frequency of *α*. Thus, we can store each proper phrase suffix along with their ranges in O(|D|) words of memory.

Now suppose that we are working on the range of the BWT of *T* corresponding to *i*-th proper phase suffix *α*. If all the occurrences of *α* in *T* are preceded by the same character `c`, then the range of the BWT of *T* associated with *α* will consist of all `c`’s. Therefore, no further computation is needed to define said range.

If the occurrences in the input of the *i*-th proper phrase suffix *α* are preceded by different characters then we make use of Lemma 2 to break the ambiguity. The order of the characters preceding *α* in *T* can be obtained from the order in which the phrases containing *α* appear in the BWT of P.

As described earlier, storing O(|P|) words can be a significant bottleneck for large repetitive datasets, and motivates the need for a BWT construction algorithm that uses less than O(|P|) words. In the next section, we introduce our recursive algorithm to address this need.

## Methods

In this section, we assume that the size of D is at least one order of magnitude smaller than the size of P—which occurs for most large, repetitive datasets—and focus on reducing the computational bottleneck that arises when the size of P increases. To accomplish this, we present a recursive solution that runs PFP on the parse, which is then used to construct the BWT of *T*. The algorithmic challenge of this approach is building BWT of *T* without access to P or its BWT. To accomplish this, we make nontrivial extensions to the algorithm that constructs the BWT from PFP [8].

### Recursive Prefix-Free Parsing

We assume that PFP was run on the input text *T* with window size *w*_1_ and integer *p*_1_. We denote the set of trigger strings defined by *w*_1_ and *p*_1_ as E_1_, and the output as P_1_ and D_1_. Next, we run PFP on the parse P_1_ with window size *w*_2_ and integer *p*_2_. We refer to running PFP on P_1_ as the *recursive step*, and denote the set of trigger strings defined by *w*_2_ and *p*_2_ as E_2_. We define the output of running PFP a second time as P_2_ and D_2_. Moreover, we denote the set of proper phrase suffixes of length greater than *w*_1_ of the phrases in D_1_ as 𝒮_1_ and, analogously, we denote the set of proper phrase suffixes of length greater than *w*_2_ of the phrases in D_2_ as 𝒮_2_. Lastly, we denote as Σ_1_ the alphabet of *D*_1_ and Σ_2_ the alphabet of *D*_2_.

As described in the previous section, the BWT ranges corresponding to proper phrase suffixes of D_1_ preceded by occurrences of the same character, can be computed just by iterating over the suffixes of D_1_ making use of Corollary 1. We note that defining the BWT of these ranges does not make use of the parse or the BWT of the parse so no advancement to `Big-BWT` is needed for these suffixes. Therefore, we focus only on the remaining proper phrase suffixes and give an equivalent to Lemma 2 that does not require direct access to P_1_ or to its BWT.

### Building the BWT from the Recursive Prefix-Free Parse

As previously mentioned, we only consider the phrases for which the `Big-BWT` algorithm needs P_1_ and/or BWTP_1_. Our goal is to compute the BWT of the input string without these. Our method for accomplishing this will rely on the following lemma.

#### Lemma 3.

*If two characters T* [*i*] *and T* [*j*] *in phrases* P_1_[*i*′] *and* P_1_[*j*′] *are followed by the same phrase suffix α* _1_, *then T* [*i*] *precedes T* [*j*] *in the BWT of T if one of the following two conditions is true: (a)* P_1_[*i*′] *and* P_1_[*j*′] *precede two different phrase suffixes α*′, ∈ *β*′ 𝒮_2_ *with α*′ ≺ *β*′; *or (b) the phrase* P_2_[*k*′] *containing* P_1_[*i*′] *precedes the phrase* P_2_[*l*′] *containing* P_1_[*j*′] *in the BWT of* P_2_.

*Proof*. We begin with proving (a). Analogously to Corollary 1 for PFP(*T*), the set of proper phrase suffixes, which we denote as 𝒮_2_, is a prefix-free set. Therefore, given that by definition *α* followed by the phrases stored in *α*′ is a prefix of *t* = *T* [*i* + 1..*n*], and *α* followed by the phrases stored in *β*′ is a prefix of *t*′ = *T* [*j* + 1..*n*], *α*′ ≺ *β*′implies *t* ≺ *t*′. Next, we prove the statement corresponding to (b). By definition, we assume that the two suffixes *t* = *T* [*i* + 1..*n*] and *t*′ = *T* [*j* + 1..*n*] start with *α* ∈ 𝒮_1_. We let *q* and *q*′ be the suffixes of P_1_ encoding the last *w*_1_ characters of those occurrences of *α* and the remainder of *t* and *t*′. Further, we let *γ* ∈ 𝒮_2_ be the prefix of *q* and *q*′. We denote *r* and *r*′ as the suffixes of P_2_ storing *q* and *q*′. Given that *α* occurs at least twice, and therefore, *γ* occurs at least twice, there exists a character where *t* and *t*′ differ. We let the first character where they differ be `c`. Given that the phrases of *D*_1_ are represented in P_1_ by their lexicographic rank. Moreover, given that the lexicographic ordering of the phrases in *D*_2_ reflects the lexicographic ordering of their expansions, and given that the elements of *D*_2_ are represented in P_2_ by their lexicographic rank, it follows from the existence of *c* that *q* ≺ *q*′ and *t* ≺ *t*′.

Using Lemma 3 we can break ambiguities for which we would need access to the BWT of P_1_ by only accessing the parse and dictionary of the recursive step, namely D_2_ and P_2_. Based on Corollary 1, Lemma 2 and Lemma 3, we can define the following data structures that we will use to compute the BWT of the input *T*.

#### Definition 1.

*We define a table* 𝒯_1_ *containing* O(|*S*_1_|) *rows and* O(1) *columns, such that for each α* ∈ 𝒮_1_, *we store in* 𝒯_1_ *its range in the BWT of T along with the colexicographic sub-range of the elements of* D_1_ *which store the occurrence of α. That is, for each c* ∈ Σ_1_ *the columns of* 𝒯_1_ *store the range of co-lexicographically sorted phrases that end in α and have c in position* |*α*| + 1 *from the end*.

Next, we define the table 𝒯_2_.

#### Definition 2.

*We define a table* 𝒯_2_ *containing* O(_2_) *rows and* O(1) *columns, such that for each α*′ ∈ 𝒮_2_, *we store in* 𝒯_2_ *the co-lexicographic range of the phrases of* D_2_ *that contain α*′ *along with the meta-characters that precede α*′ *in P*_1_.

Lastly, we define the grid 𝒢_2_.

#### Definition 3.

*We define the grid* 𝒢_2_ *containing* O(|*P*_2_|) *rows and* O(|*D*_2_|) *columns, such that for each element* ℓ *of* D_2_, 𝒢_2_ *stores the positions in the BWT of* P_2_ *where* ℓ *appears*.

The data structures just defined allow us to compute the characters of the BWT of *T* that require Lemma 3 through the following algorithm. We assume that we computed the BWT for all proper phrases suffixes in 𝒮_1_ up to *α*. From 𝒯_1_, we know that *α* is preceded in *T* by the characters `c`_1_ and `c`_2_, and that `c`_1_ is stored in phrase *p*_1_ ∈ D_1_, and that `c`_2_ is stored in phrase *p*_2_ ∈ D_2_. In other words, we cannot break the ambiguity using only 𝒯_1_. We now list, following their lexicographic order, all proper phrase suffixes in *S*_2_ that are preceded in *P*_1_ by *p*_1_, *p*_2_ or both using 𝒯_2_. First, we assume that we are in case (a) and thus, we find (using 𝒯_2_) the phrase is preceded by *p*_1_, say *α*′. Using Lemma 3, we see that the corresponding character can be defined without any further computation.

Next, we assume that we find (using 𝒯_2_) the phrase that is preceded by both *p*_1_ and *p*_2_, say *β*′. It follows from Lemma 3 that in order to find the corresponding characters in BWT_*T*_, we need to consider the relative order of the occurrences of the phrases storing *β*′ and preceded by *p*_1_ and *p*_2_ in the BWT of P_2_. We can find the co-lexicographic range of elements of D_2_ ending in *β*′, and for each occurrence of those phrases in 𝒢_2_, we can define the BWT_*T*_.

Thus, we summarize our result with the following theorem, which directly follows from Corollary 1, Lemma 2 and Lemma 3 using the data structures 𝒯_1_, 𝒯_2_. and 𝒢_2_

#### Theorem 1.

*Given a string T, the dictionary* D_1_ *and the parse* P_1_ *obtained by running PFP on T, and the dictionary* D_2_ *and the parse* P_2_ *obtained by running PFP on* P_1_, *we can compute the BWT*_*T*_ *from* D_1_, D_2_ *and* P_2_ *using* O(|D_1_| + |D_2_| + |P_2_|) *workspace*.

## Experiments and Results

### Experimental Set-up

We implemented `r-pfbwt` in ISO C++ 2020 and measured the performance using a real world genomic dataset. It consists of 12 sets of variants of human chromosome 19 (`chr19`), containing 200, 400, 600, 800, 1000, 1200, 1400, 1600, 1800, 2000, 2200 and 2400 distinct individuals. Each individual consists of 2 sequences (haplotypes). Hence, we evaluated our method on up to 4800 sequences. Each collection is a superset of the previous one. The smallest of these datasets (`chr19.200`) has size 22GB and the largest (`chr19.2400`) has size 264GB with each data point increasing in size by 22GB. Table 1 reports input size, the size of the parse and dictionary of the input, and the size of the parse and dictionary of the recursive step. We compared `r-pfbwt` to `Big-BWT` given that this out-performed all competing methods to build the RLBWT as shown in [9]. The running time and the maximal resident set size were recorded with the Unix utility `/usr/bin/time`. Given that both methods require building PFP from the input the time required for its construction are omitted from the total.

**Table 1:** In order to illustrate the advantage of our recursive algorithm, we illustrate the size of the input, the size of the dictionary and parse from prefix-free parsing of the input sequences, and the size of the dictionary and parse from prefix-free parsing PFP(*T*). All sizes are shown in gigabytes.

| Haplotypes | Input Size | $\vert P_1\vert $ | $\vert D_1\vert $ | $\vert D_1\vert + \vert P_1\vert $ | $\vert P_2\vert $ | $\vert D_2\vert $ | $\vert D_2\vert + \vert P_2\vert $ |
| --- | --- | --- | --- | --- | --- | --- | --- |
| 200 | 22.08 | 0.96 | 0.16 | 1.11 | 0.08 | 0.05 | 0.13 |
| 400 | 44.11 | 1.91 | 0.23 | 2.14 | 0.16 | 0.09 | 0.25 |
| 600 | 66.13 | 2.86 | 0.27 | 3.13 | 0.24 | 0.12 | 0.36 |
| 800 | 88.16 | 3.82 | 0.32 | 4.14 | 0.32 | 0.15 | 0.48 |
| 1000 | 110.18 | 4.77 | 0.36 | 5.13 | 0.73 | 0.16 | 0.89 |
| 1200 | 132.21 | 5.72 | 0.40 | 6.13 | 0.49 | 0.22 | 0.71 |
| 1400 | 154.24 | 6.68 | 0.43 | 7.11 | 0.57 | 0.25 | 0.82 |
| 1600 | 176.26 | 7.63 | 0.46 | 8.09 | 0.65 | 0.27 | 0.92 |
| 1800 | 198.29 | 8.59 | 0.48 | 9.07 | 0.73 | 0.29 | 1.02 |
| 2000 | 220.31 | 9.54 | 0.51 | 10.05 | 0.81 | 0.31 | 1.12 |
| 2200 | 242.34 | 10.49 | 0.54 | 11.03 | 0.89 | 0.34 | 1.22 |
| 2400 | 264.36 | 11.45 | 0.56 | 12.00 | 0.97 | 0.35 | 1.32 |

## Results

On Chromosomes 19 `r-pfbwt` required less memory than `Big-BWT` and was able to complete the execution in less time. In fact, on 2400 chromosomes, `r-pfbwt` was 2 times faster and required 2.5 times less memory. Moreover, the empirical results demonstrate that the performance gains increase as the data gets larger. `r-pfbwt` was 1.1 times faster and required 1.2 times less memory on 1000 chromosomes, 2.0 times faster and 2.3 times less memory on 2000 chromosomes and 2.2 times faster and 2.3 times less memory on 2400 chromosomes.

## Conclusions

When indexing large repetitive datasets using PFP the size of the parse P quickly becomes the computational bottleneck requiring significant amounts of memory. In this work we address the challenge of reducing the total memory required by running PFP again on P. We show that is possible to use only memory proportional to the size of the dictionary obtained by running PFP on the input text and to the size of the dictionary and parse obtained with the *recursive step*. In this work we first show the correctness of our approach and, through experiments on real world datasets, we show its effectiveness in reducing the memory required to build the BWT of the input. Moreover, we implemented our method introducing more efficient parallelization compared to `Big-BWT`, reducing the total wall clock time required. Lastly, it is worth noting that the technique presented in this paper can be applied to reduce the memory footprint for many of the applications of PFP proposed in the last few years.

**Figure 1:**
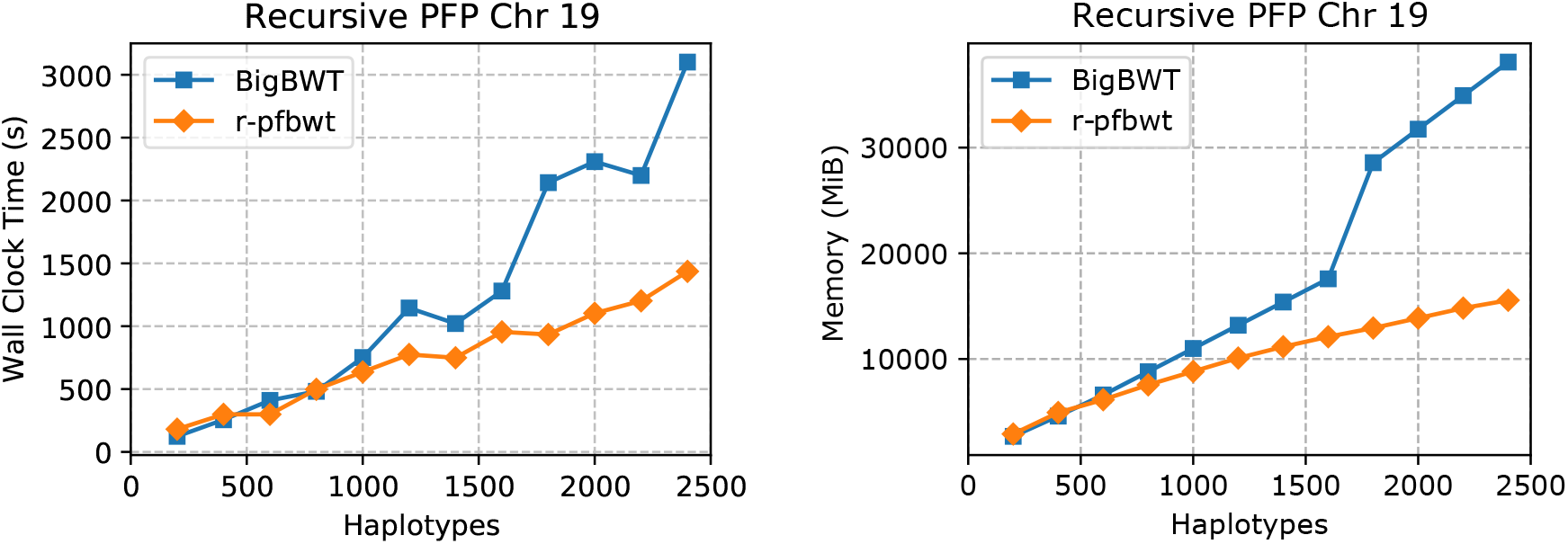
Chromosomes 19 construction Wall Clock Time in seconds (**left**) and peak memory in MiB (**right**).

## Footnotes

1 We note that this definition is equivalent to original definition that considers the string *T* ^*′′*^ = *S*#^*w*^ to be circular.

## References

[1] Michael Burrows and David Wheeler, “A block-sorting lossless data compression algorithm,” in Digital SRC Research Report. Citeseer, 1994.

[2] H. Li, “Aligning sequence reads, clone sequences and assembly contigs with BWA-MEM,” arXiv, p. http://arxiv.org/abs/1303.3997, 2013.

[3] B. Langmead and S.L. Salzberg, “Fast gapped-read alignment with Bowtie 2,” Nature Methods, vol. 9, no. 4, pp. 357–359, 2012.

[4] H. Li, “Fast construction of FM-index for long sequence reads,” Bioinformatics, vol. 30, no. 22, pp. 3274–3275, 2014.

[5] J. Sirén, “Burrows-wheeler transform for terabases,” in Proc. of IEEE Data Compression Conference (DCC), 2016, pp. 211–220.

[6] C. Boucher, T. Gagie, A. Kuhnle, B. Langmead, G. Manzini, and T. Mun, “Prefix-free parsing for building big BWTs,” Algorithms in Molecular Biology, vol. 14, no. 1, pp. 13:1–13:15, 2019.

[7] U. Manber and G. W. Myers, “Suffix arrays: a new method for on-line string searches,” SIAM Journal on Computing, vol. 22, no. 5, pp. 935–948, 1993.

[8] C. Boucher, T. Gagie, A. Kuhnle, B. Langmead, G. Manzini, and T. Mun, “Prefix-free parsing for building big BWTs,” Algorithms for Molecular Biology, vol. 14, no. 1, pp. 13, Dec. 2019.

[9] C. Boucher, T. Gagie, A. Kuhnle, and G. Manzini, “Prefix-free parsing for building big BWTs,” in Proc. of Workshop on Algorithms in Bioinformatics WABI, 2018, pp. 2:1–2:16.

